# Regional Mutational Signature Activities in Cancer Genomes

**DOI:** 10.1101/2022.01.23.477261

**Authors:** Caitlin Timmons, Quaid Morris, Caitlin F. Harrigan

## Abstract

Cancer genomes harbor a catalog of somatic mutations. The type and genomic context of these mutations depend on their causes, and allow their attribution to particular mutational signatures. Previous work has shown that mutational signature activities change over the course of tumor development, but investigations of genomic region variability in mutational signatures have been limited. Here, we expand upon this work by constructing regional profiles of mutational signature activities over 2,203 whole genomes across 25 tumor types, using data aggregated by the Pan-Cancer Analysis of Whole Genomes (PCAWG) consortium. We present GenomeTrackSig as an extension to the TrackSig R package to construct regional signature profiles using optimal segmentation and the expectation-maximization (EM) algorithm. We find that 426 genomes from 20 tumor types display at least one change in mutational signature activities (changepoint), and 257 genomes contain at least one of 54 recurrent changepoints shared by seven or more genomes of the same tumor type. Five recurrent changepoint locations are shared by multiple tumor types. Within these regions, the particular signature changes are often consistent across samples of the same type and some, but not all, are characterized by signatures associated with subclonal expansion. The changepoints we found cannot strictly be explained by gene density, mutation density, or cell-of-origin chromatin state. We hypothesize that they reflect a confluence of factors including evolutionary timing of mutational processes, regional differences in somatic mutation rate, large-scale changes in chromatin state that may be tissue type-specific, and changes in chromatin accessibility during subclonal expansion. These results provide insight into the regional effects of DNA damage and repair processes, and may help us localize genomic and epigenomic changes that occur during cancer development.

## Introduction

Cancer is a disease that develops over a lifetime through a series of somatic mutations. The vast majority of these mutations are passenger mutations, which have little effect on fitness (Dietlein et al., 2020, Martincorena et al., 2017). Multiple mutagenic processes operate in a cancer throughout its development, which in turn determine the nature of the collection of somatic mutations that accumulate. Different processes give rise to distinct mutational patterns, called *mutational signatures* (Alexandrov et al., 2020, Koh et al., 2021). In nearly all cancers, multiple mutagenic processes contribute somatic mutations to individual cancer genomes. The relative contributions of these different processes can be quantified by algorithms that assign an activity (or exposure) to each detected mutational signature (Alexandrov et al., 2020). Recent work (Dentro et al., 2021, Gerstung et al., 2020, Rubanova et al., 2020) has revealed that i) these activities vary during cancer development (i.e., tumor progression) in a cancer-type-specific way and ii) most mutational signatures are primarily active only in either early- or late-tumor progression (Dentro et al., 2021, Gerstung et al., 2020). Computational methods, like TrackSig, have been developed to reconstruct the evolutionary trajectories of mutational signature activities in individual samples (Rubanova et al., 2020, Harrigan et al., 2020). As such, mutational signature analysis is a compelling tool in our search for understanding of the processes shaping tumor formation and development.

In addition to evolutionary timing of signature activity, somatic mutation rate and mutational signatures have been found to be influenced by genomic and epigenomic factors that exert their effects on the scale of several nucleotides up to a megabase. These factors include sequence context, nucleosome positioning, chromatin state, and replication timing (e.g. Gonzalez-Perez et al. (2019), Haradhvala et al. (2016), Hodgkinson et al. (2012), Lawrence et al. (2013), Polak et al. (2014, 2015), Schuster-Böckler and Lehner (2012), Seplyarskiy and Sunyaev (2021), Supek and Lehner (2019), Vöhringer et al. (2021), Yaacov et al. (2021)). For example, late-replicating and heterochromatic regions accumulate more mutations than early-replicating and euchromatic regions (Supek and Lehner, 2019, 2015, Zheng et al., 2014). This results in somatic mutation accumulation patterns that can be attributed to chromatin state in the cell of origin. For example, in mismatch repair (MMR)-proficient tumors, the rate of CpG>TpG mutations may correlate with replication timing since MMR corrects mispairings due to 5mC deamination more efficiently in early-replicating areas (Gonzalez-Perez et al., 2019, Supek and Lehner, 2019).

These mutation-associated genomic features vary on scales of 1 Mb or less, so analyses of their effects on mutational signature activity are typically confined to particular loci. To our knowledge, larger scale relationships between chromosomal location and mutational signature activity have not been thoroughly investigated. In particular, if there exists common evolutionary events that dictate the action of mutational processes in cancers of the same type, we can expect to observe consistent changes to mutational signature activity across genomes.

Also, changepoint identification yields both biological and potential clinical insights. Many mutational signatures are associated with defects in DNA repair processes; these processes can also be inhibited by clinical treatments (Gavande et al., 2016). Changepoint analysis may help us understand how these treatments’ effects vary across the genome. With GenomeTrackSig, we can also explore how changes in genomic features such as copy number or chromatin state that occur during tumorigenesis affect the activity of mutational processes on a genome-wide scale. The changepoints we identify can also lend insight into intra-tumor heterogeneity, which presents challenges from a therapeutic perspective. Changes in signature activity may mark parts of the genome more susceptible to particular mutational processes early vs. late in tumor development. Further, changepoints that recur across multiple samples and tumor types may further our understanding of tumorigenesis by identifying regional DNA damages or repair deficiencies that are characteristic of a tumor type or a group of tumor types.

To examine the occurrence of mutational signature activity changes across the genome, we analyzed somatic mutations of 2,203 whole genomes across 25 tumor types from the Pan-Cancer Analysis of Whole Genomes (PCAWG) consortium (Campbell et al., 2020). Using GenomeTrackSig, an extension of TrackSig (Rubanova et al., 2020), we identified genomic regions that recurrently exhibit mutational signature changes. These regions frequently showed activity changes in signatures known to be associated with evolutionary timing. Some of these regions were shared among several tissue types, while others appeared to be exclusive to a single tissue.

## Methods

GenomeTrackSig, an extension of the TrackSig algorithm (Rubanova et al., 2020, Harrigan et al., 2020), is designed to identify changes in mutation signature activity across the genome. TrackSig was designed to detect changes in mutational signature activity across mutations ordered by pseudotime. TrackSig segments a set of mutations, sorted by inferred cancer cell frequency to determine changes in mutational signature activity. We extended TrackSig such that instead of considering evolutionary timing of mutations, we can detect changes in signature activity across chromosomal coordinates. An overview of our methodology is shown in Figure 1. In accordance with the original TrackSig algorithm, we represent a sample containing N mutations as an *NXK* matrix where each mutation is given as a binary vector over *K* mutation types. This matrix is in turn represented as a mixture of signature multinomials, in which the mixture coefficients are interpreted as the signature activities, or the probability of a given mutation *n* to be assigned to a given signature, *s*_*i*_. The probability that mutation n was generated by a signature *s*_*i*_ is given by the *k*^*th*^ component of *s*_*i*_ raised to the power of the *k*^*th*^ component of the binary mutation vector, multiplied across *K* components. We use the EM algorithm (Moon, 1996) to fit the mixture coefficients in each segment.

**Fig. 1.**
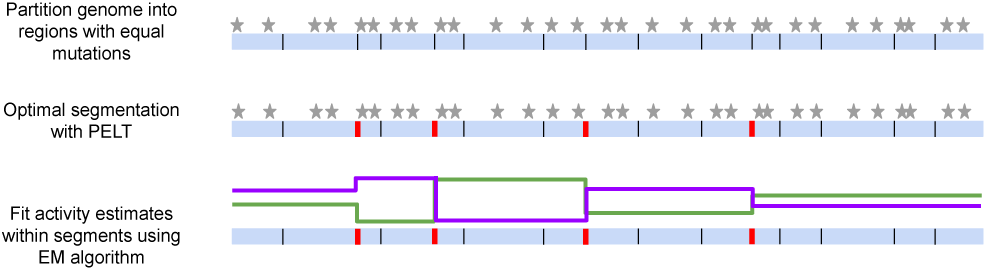
Overview of GenomeTrackSig algorithm for profiling mutational signature activities across the genome. Stars indicate mutations (top); red bars indicate changepoints (middle); green and purple lines indicate estimated mutation signature activities; y-axis indicates exposure level (bottom). GenomeTrackSig requires at least 100 mutations per segment, fewer mutations are shown here for illustrative purposes.

We identify changepoints using the Pruned Exact Linear Time (PELT) segmentation algorithm (Killick et al., 2012). PELT uses dynamic programming and branch and bound search to perform optimal segmentation which is efficient in the number of mutations. At each iteration, PELT considers adding a new changepoint out of the set of available regions and scores the changepoint by refitting the activities in each region. Over-segmentation is penalized using the Bayesian Information Criterion (BIC). Previous work by Alexandrov et al. (2020) characterized canonical signatures active within the PCAWG data. These signatured serve as the set of reference signatures for our analyses. We fit activity estimates to all signatures which have been previously determined (Alexandrov et al., 2020) to be active in a given tumor type. GenomeTrackSig extends the capabilities of the original TrackSig algorithm to explore how mutational signature activities vary across chromosomal regions in a variety of tumors.

By default, we partition the set of mutations in a genome into bins of 100 mutations each, where the size of the chromosome region (i.e., number of basepairs) spanned by a bin will vary depending on mutation density. We require each segment in the segmentation solution to include at least one bin. GenomeTrackSig includes flexible options for analyzing samples, either by partitioning and estimating signature activities across an entire genome or on each individual chromosome (i.e., a changepoint is always placed at chromosomal boundaries). The genome-wide approach can capture signature activities that span multiple chromosomes thereby potentially reducing noise in activity estimates at chromosomal ends. On the other hand, the chromosome-wise approach may be more sensitive to activity changes within a chromosome and substantially reduces runtime on samples with many mutations because each chromosome can be segmented in parallel (Figure S1). In general, the chromosome-wise approach is preferred, but we use the genome-wide strategy for samples (and cancers) where chromosomes have less than 100 mutations, as we use this as a minimum segment size to ensure accurate activity estimates. Similar to TrackSig, we find that GenomeTrackSig is relatively insensitive to the choice of bin size when determining changepoint placement (Figure S2).

## Results

### A. Mutational signature activity is not constant across the genome

We constructed genome-wide signature activity profiles for 2,203 tumors across 25 tumor types (Biliary-AdenoCA, Bladder-TCC, Bone-Osteosarc, Breast-AdenoCA, Cervix, CNS-GBM, Colorect-AdenoCA, Eso-AdenoCA, Head-SCC, Kidney-ChRCC, Kidney-RCC, Liver-HCC, Lung-AdenoCA, Lung-SCC, Lymph-BNHL, Lymph-CLL, Melanoma, Myeloid-MPN, Ovary-AdenoCA, Panc-AdenoCA, Panc-Endocrine, Prost-AdenoCA, Stomach-AdenoCA, Thy-AdenoCA, Uterus-AdenoCA) (Table S1). The number and width of bins in each tumor varied from 23-2,895 and 1 Mb to 250 Mb, respectively, based on the number and distribution of mutations in the tumor (Figure S3, S4). In total, we observed changepoints in 426 of 2,203 samples analyzed. We found no genomic changepoints in five types of tumor: Liver-HCC (N=326), Ovary-AdenoCA (N=113), Biliary-AdenoCA (N=29), Panc-Endocrine (N=76), and Myeloid-MPN (N=31). A lack of change in signature activity could reflect an even mutational composition across the genome, insufficient number of mutations to characterize a change in signature activity, a lack of large-scale variation in chromatin accessibility, or a lack of substantial changes in mutational composition over tumor development. In the remaining 20 tumor types, at least one sample exhibited changes in signature activities across the genome, indicating that large-scale shifts in signature activities depending on chromosomal location are a common but not ubiquitous phenomenon (Table 1).

**Table 1.**
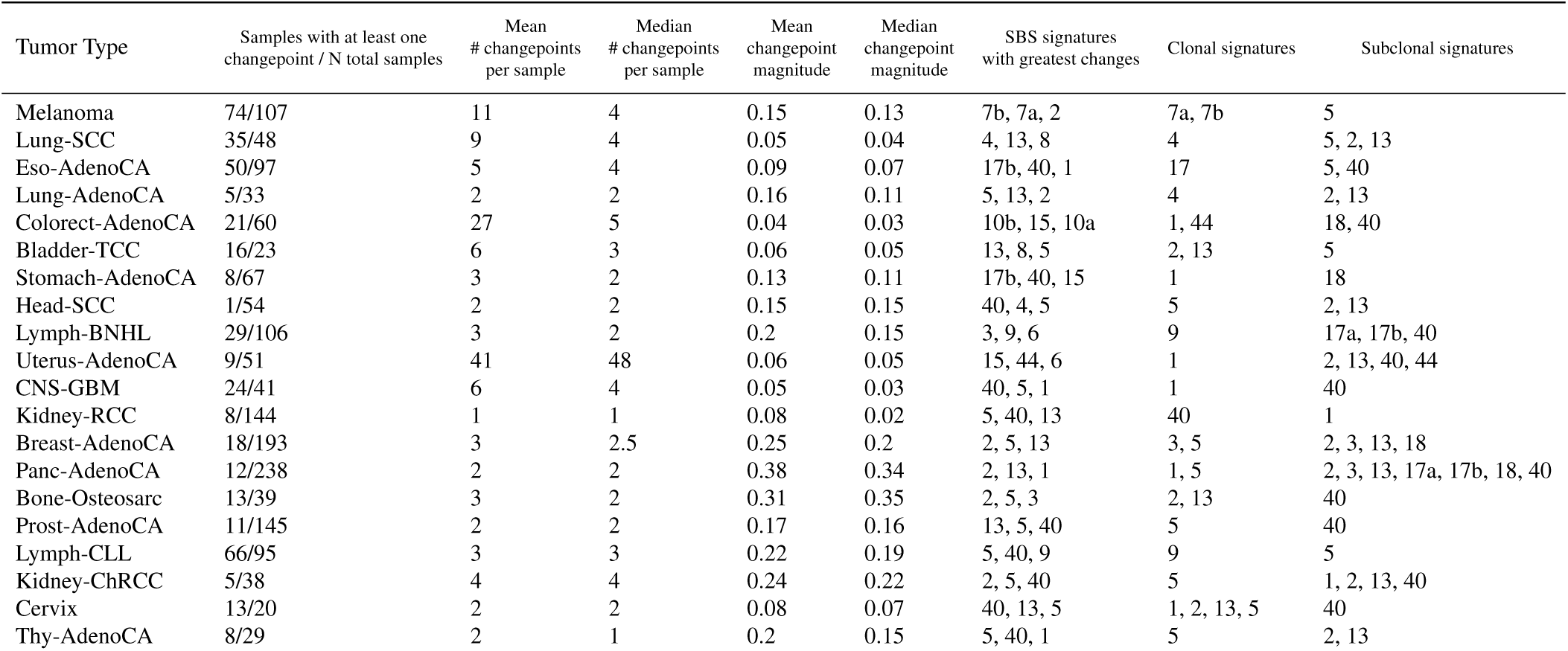
Overview of changepoints discovered across 20 cancer types. Magnitude of each changepoint is measured as the cosine distance of the signature activity vector on either side of the changepoint. For each cancer type containing changepoints, the three signatures with the greatest absolute value of activity changes across all changepoints in that cancer are listed. Signatures are indicated as “clonal” or “subclonal” respectively based on where their activity is highest, as described by Dentro et al. (2021).

| Tumor Type | Samples with at least one changepoint / N total samples | Mean # changepoints per sample | Median # changepoints per sample | Mean changepoint magnitude | Median changepoint magnitude | SBS signatures with greatest changes | Clonal signatures | Subclonal signatures |
| --- | --- | --- | --- | --- | --- | --- | --- | --- |
| Melanoma | 74/107 | 11 | 4 | 0.15 | 0.13 | 7b, 7a, 2 | 7a, 7b | 5 |
| Lung-SCC | 35/48 | 9 | 4 | 0.05 | 0.04 | 4, 13, 8 | 4 | 5, 2, 13 |
| Eso-AdenoCA | 50/97 | 5 | 4 | 0.09 | 0.07 | 17b, 40, 1 | 17 | 5, 40 |
| Lung-AdenoCA | 5/33 | 2 | 2 | 0.16 | 0.11 | 5, 13, 2 | 4 | 2, 13 |
| Colorect-AdenoCA | 21/60 | 27 | 5 | 0.04 | 0.03 | 10b, 15, 10a | 1, 44 | 18, 40 |
| Bladder-TCC | 16/23 | 6 | 3 | 0.06 | 0.05 | 13, 8, 5 | 2, 13 | 5 |
| Stomach-AdenoCA | 8/67 | 3 | 2 | 0.13 | 0.11 | 17b, 40, 15 | 1 | 18 |
| Head-SCC | 1/54 | 2 | 2 | 0.15 | 0.15 | 40, 4, 5 | 5 | 2, 13 |
| Lymph-BNHL | 29/106 | 3 | 2 | 0.2 | 0.15 | 3, 9, 6 | 9 | 17a, 17b, 40 |
| Uterus-AdenoCA | 9/51 | 41 | 48 | 0.06 | 0.05 | 15, 44, 6 | 1 | 2, 13, 40, 44 |
| CNS-GBM | 24/41 | 6 | 4 | 0.05 | 0.03 | 40, 5, 1 | 1 | 40 |
| Kidney-RCC | 8/144 | 1 | 1 | 0.08 | 0.02 | 5, 40, 13 | 40 | 1 |
| Breast-AdenoCA | 18/193 | 3 | 2.5 | 0.25 | 0.2 | 2, 5, 13 | 3, 5 | 2, 3, 13, 18 |
| Panc-AdenoCA | 12/238 | 2 | 2 | 0.38 | 0.34 | 2, 13, 1 | 1, 5 | 2, 3, 13, 17a, 17b, 18, 40 |
| Bone-Osteosarc | 13/39 | 3 | 2 | 0.31 | 0.35 | 2, 5, 3 | 2, 13 | 40 |
| Prost-AdenoCA | 11/145 | 2 | 2 | 0.17 | 0.16 | 13, 5, 40 | 5 | 40 |
| Lymph-CLL | 66/95 | 3 | 3 | 0.22 | 0.19 | 5, 40, 9 | 9 | 5 |
| Kidney-ChRCC | 5/38 | 4 | 4 | 0.24 | 0.22 | 2, 5, 40 | 5 | 1, 2, 13, 40 |
| Cervix | 13/20 | 2 | 2 | 0.08 | 0.07 | 40, 13, 5 | 1, 2, 13, 5 | 40 |
| Thy-AdenoCA | 8/29 | 2 | 1 | 0.2 | 0.15 | 5, 40, 1 | 5 | 2, 13 |

### B. Some changepoints are shared across tissue type

We find that changepoint regions are often shared—both across multiple cancers from the same tissue and across multiple tissues. To determine which changepoints within samples from the same tumor type overlap, we fit a kernel density estimate across the vector of genomic locations covered by each changepoint and constructed a possible range for that changepoint location, which contains the changepoint locus identified by GenomeTrackSig, +/- one standard deviation of the density function. We then overlaid all changepoint ranges from that tumor type and counted how many changepoints fell within a sliding window across the genome. We then deemed a “recurrent changepoint region” for each tumor type as the center of the region where at least seven samples shared an overlapping changepoint; i.e. the window within a region in which the maximum number of changepoints fell. This threshold corresponded to an elbow (Figure S5).

Of the 20 tumor types that show any changes in signature activity across their genomes, eight contain recurrent changepoint regions (Figure 2). Interestingly, the changes in signature activity at these regions are often highly similar across samples of the same tissue type, both in direction and magnitude. For example, all 55 melanoma samples that have a changepoint overlapping with the chromosome 1:47,000,000-1:51,000,000 region show a decrease in SBS7b activity and increase in SBS7a activity in the direction of the numbering (Figure 3). In Uterus-AdenoCA samples, the signatures which changed most at every recurrent change-point are associated with either defective DNA mismatch repair (SBS15) or polymerase epsilon mutations (SBS10a, SBS10b) (Figure 2). In other cancers, the most affected signatures are highly variable from changepoint-to-changepoint: seven different signatures are represented among the most substantial activity changes at recurrent changepoints in Eso-AdenoCA samples, including SBS1, SBS2, SBS3, SBS5, SBS17a, SBS17b, and SBS40 (Figure 2).

**Fig. 2.**
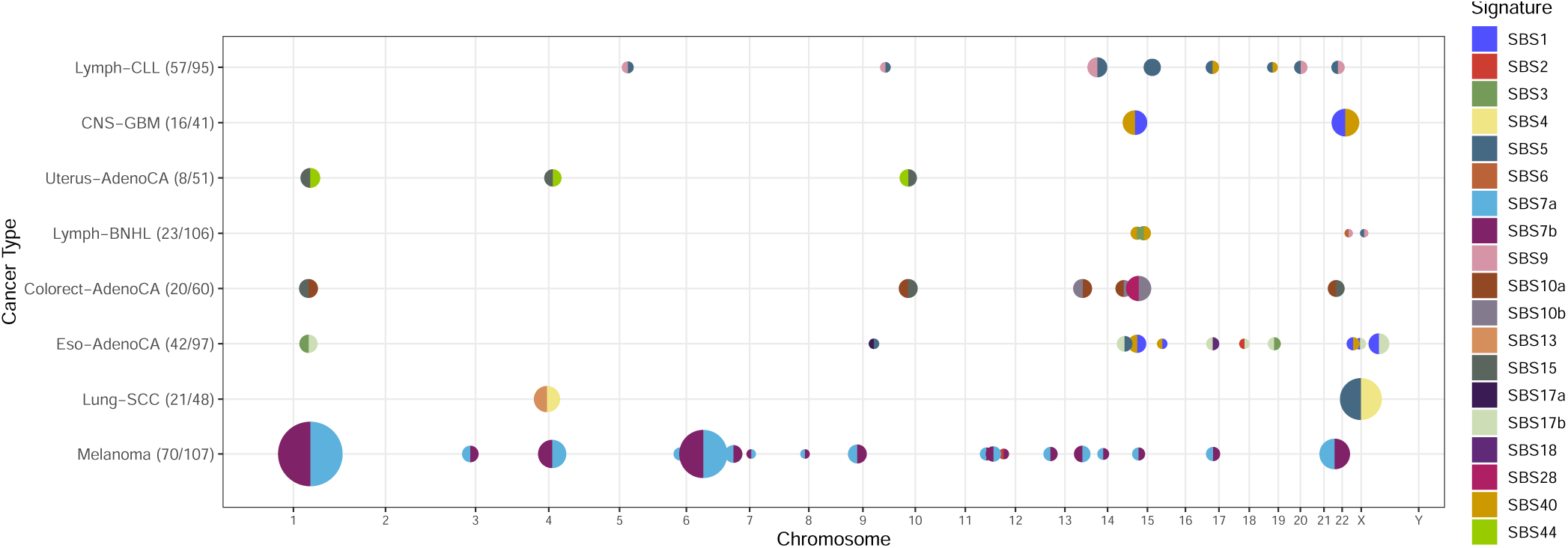
Recurrent changepoint regions across eight tumor types. 54 genomic regions include a changepoint found in seven or more samples for a given tumor type. Each of these recurrent changepoint regions are indicated by a point, where COSMIC V3 signatures Alexandrov et al. (2020), Tate et al. (2019) are indicated by color. Changepoints are summarized by the signature whose activity decreases the most on average (left half) and the signature whose activity increases the most on average (right half). Proportion of samples exhibiting a changepoint is encoded by size, and the number of samples affected is indicated in the tumor type label (n/N). Changepoints that appear in fewer than seven samples are omitted.

**Fig. 3.**
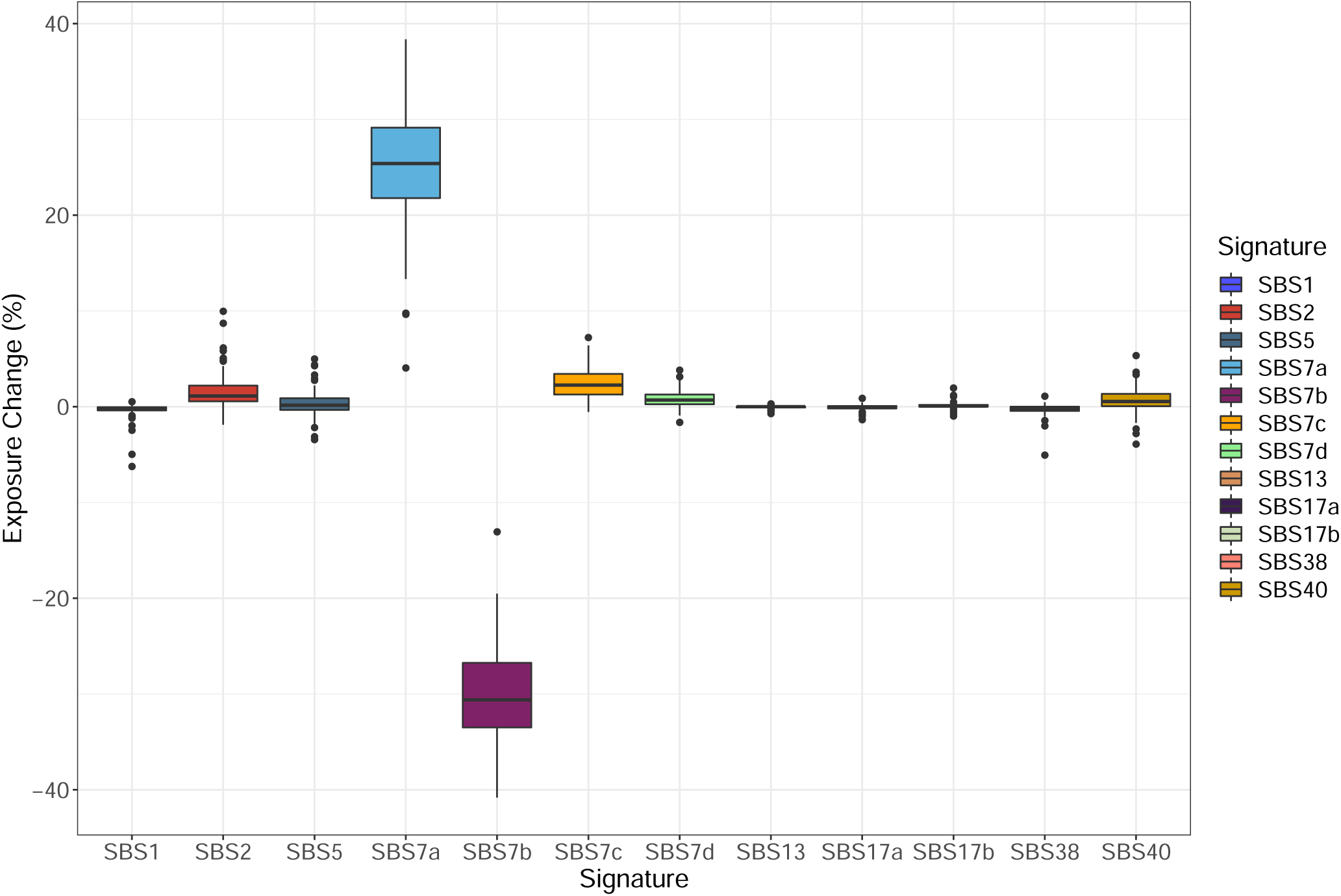
Similar signature changes at the 1:47,000,000-1:51,000,000 region in 55/107 melanoma samples. Each boxplot shows the distribution of activity changes for a particular signature at the recurrent changepoint region 1:47,000,000-1:51,000,000.

We also find that five recurrent changepoints are shared by two tumor types (Figure 2). The shared recurrent changepoint at 1:47,000,000-1:51,000,000 appears most frequently, in 55 Melanoma and eight Uterus-AdenoCA samples. Another recurrent changepoint region at 1:42,000,000-1:47,000,000 is shared by 14 Eso-AdenoCA and nine Colorect-AdenoCA samples.

Despite the consistency of changepoint locations across multiple samples and tissues, it is unlikely that all these change-points result from mutations in specific genes near the changepoint region. For one, changepoint regions span multiple megabases, so it is difficult to attribute the change-point to mutations in any one particular gene. That said, some changepoint regions contain genes that may play a role in tumorigenesis such as Lymph-CLL changepoint regions encompassing CBL proto-oncogene B and NRP2 (Goel et al., 2012, Liyasova et al., 2015), and a changepoint region shared by Lung-SCC and Bladder-TCC which encompasses the DDX17 gene (Wu, 2020). However, we were unable to detect any clear pattern of gene function at changepoint regions.

### C. Many changepoints are characterized by signatures associated with subclonal expansion

In certain cancers, we find that recurrent changepoints show a shift in activity from signatures associated with early cancer development to signatures associated with subclonal expansion. For instance, at five of eight recurrent changepoints in Lymph-CLL samples, the most dramatic signature activity changes were increases in SBS9 and decreases in SBS5, or vice versa. We validated that these changepoints indeed reflect a shift between early and late signatures by constructing evolutionary trajectories for 90 Lymph-CLL samples using TrackSig (Rubanova et al., 2020). Figure 4 shows the distribution of activity changes from early to late development across all samples, in which we observe a definitive decrease in SBS9, and increase in SBS5 activity. In other cancers, signatures which account for the most dramatic activity changes at recurrent changepoint regions do not show a clear timing association. For example, SBS7a and SBS7b change most dramatically across the genome in melanoma samples (Figure 2), yet do not show as strong of an association with evolutionary timing (from early-occurring to late-occuring mutations, SBS7a activity decreases by 8.7% +/- 12.2% on average and SBS7b activity increases by 1.7% +/- 9.1% on average).

**Fig. 4.**
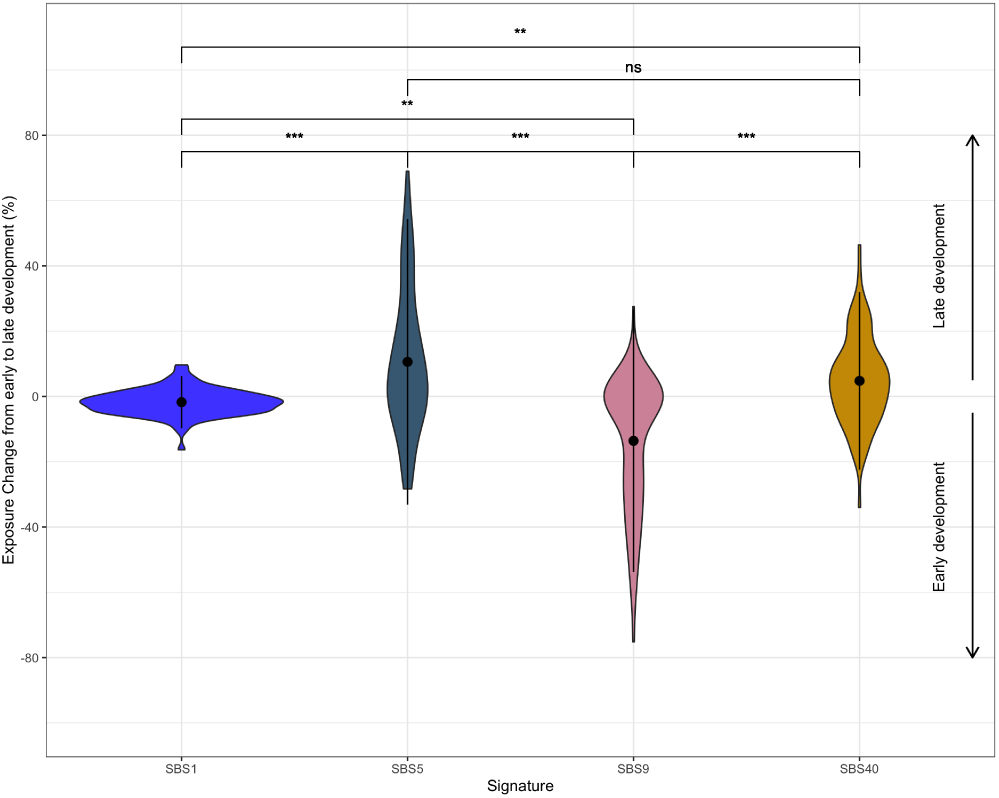
Activity change in four SBS signatures during cancer evolution in 90 Lymph-CLLs. Violin plots showing the distribution of signature activity changes (y-axis) from early to late stages of cancer development constructed using TrackSig (Dentro et al., 2021, Rubanova et al., 2020, Harrigan et al., 2020) for 90 chronic lymphocytic leukemia samples. Dot indicates means, vertical line spans +/- one standard deviation. From early to late development, SBS1 changes on average by -1.8% +/- 4.0%, SBS5 by 10.6% +/- 21.9%, SBS9 by -13.6% +/- 20.1%, and SBS40 by 4.7% +/- 13.6%. Brackets display p-values for Kolmogorov-Smirnov tests between activity change distributions, adjusted for multiple comparisons with Bonferroni corrections. * indicates adj. p < 1e-2, ** indicates adj. p < 1e-5, *** indicates adj. p < 1e-8.

Chronic lymphocytic leukemias (Lymph-CLL) tend to exhibit high SBS9 activity early in development, and show decreasing SBS9 and increasing SBS5 and SBS40 activities as subclones form (Figure 4), (Dentro et al., 2021). SBS9 activity is attributed to mutagenesis induced via DNA polymerase eta. These mutations can occur in healthy lymphoid cells as part of a somatic hypermutation process, which introduces mutations to antibody-coding sequence in order to generate sequence variability and produce antibodies with higher specificity (Seki et al., 2005). SBS9 activity is elevated in Lymph-CLL samples that possess immunoglobulin gene hypermutation (Alexandrov et al., 2020, Gerstung et al., 2020), a mutational process that typically occurs early in tumor development (Dentro et al., 2021, Seifert et al., 2012). Interestingly, changes in signature activity were distributed at many loci in Lymph-CLL samples, whereas in B-cell non-Hodgkin lymphoma (Lymph-BNHL), which also undergoes polymerase eta dependent somatic hypermutation, changes in signature activity were concentrated on chromosome 14, close to the immunoglobulin gene hypermutation (IGH) locus. While the etiologies of SBS5 and SBS40 are unknown, their activity correlates with patient age, and SBS5 has been associated with proliferation (Alexandrov et al., 2020, Franco et al., 2019).

Lymph-CLL signature activity has previously been seen to be associated with evolutionary timing. Similar cancers also evolve in consistent patterns (Dentro et al., 2021, Rubanova et al., 2020, Campbell et al., 2020). In keeping with this, we observe recurrent changepoints that may be indicative of important evolutionary (and shared) changes that occur over the progression of these tumors, such as timing-dependent changes in chromatin state at these regions. Regional mutation rates and mutational signatures are influenced by chromatin state, and under normal cell growth conditions we would expect to see higher mutation rates in heterochromatic than euchromatic regions (Vöhringer et al., 2021, Yaacov et al., 2021). However, the compounding DNA damage and dysfunction in DNA repair that occurs as cancers develop could induce changes in chromatin state, making regions vulnerable to mutations late in development that were not vulnerable during early cancer formation. As such, if the chromatin state at a particular region changes during tumor evolution, this may also manifest as a regional change in signature activities.

### D. Changepoints are not associated with any single phenomenon

We examined the correlation between changepoint locations and changes in gene density, mutation density, and when available, tissue-specific measures of chromatin accessibility (Polak et al., 2015) to investigate what underlying genomic or epigenomic features could account for the observed distribution of changepoints. Given that tissue-specific or cancer-specific DNAse-Seq data is not available for all tumor types in our dataset, we cannot draw broad conclusions about the relationship between local changes in chromatin accessibility and changepoint placement. However, it is interesting to note that most melanoma samples display low correlation between DNAse-I accessibility index and changepoint occurrence (mean Pearson correlation coefficient, R=0.12), thus it does not seem that local chromatin state changes alone can explain changes in signature activity (Figure 5). Changes in gene density or mutation density alone also fail to explain the distribution of observed changepoints. For instance, the mean correlations between changepoint occurrence and gene and mutation density across 74 melanoma samples are R=0.13 and R=0.11, respectively (Figure 3). Changepoints occur in both gene and mutation-dense and gene and mutation-poor regions, changes in both variables are often unaccompanied by changepoints, and we could observe no obvious biased placement of changepoint with respect to these variables. Furthermore, recurrent changepoints occur across multiple samples and cancer types, where these features vary.

**Fig. 5.**
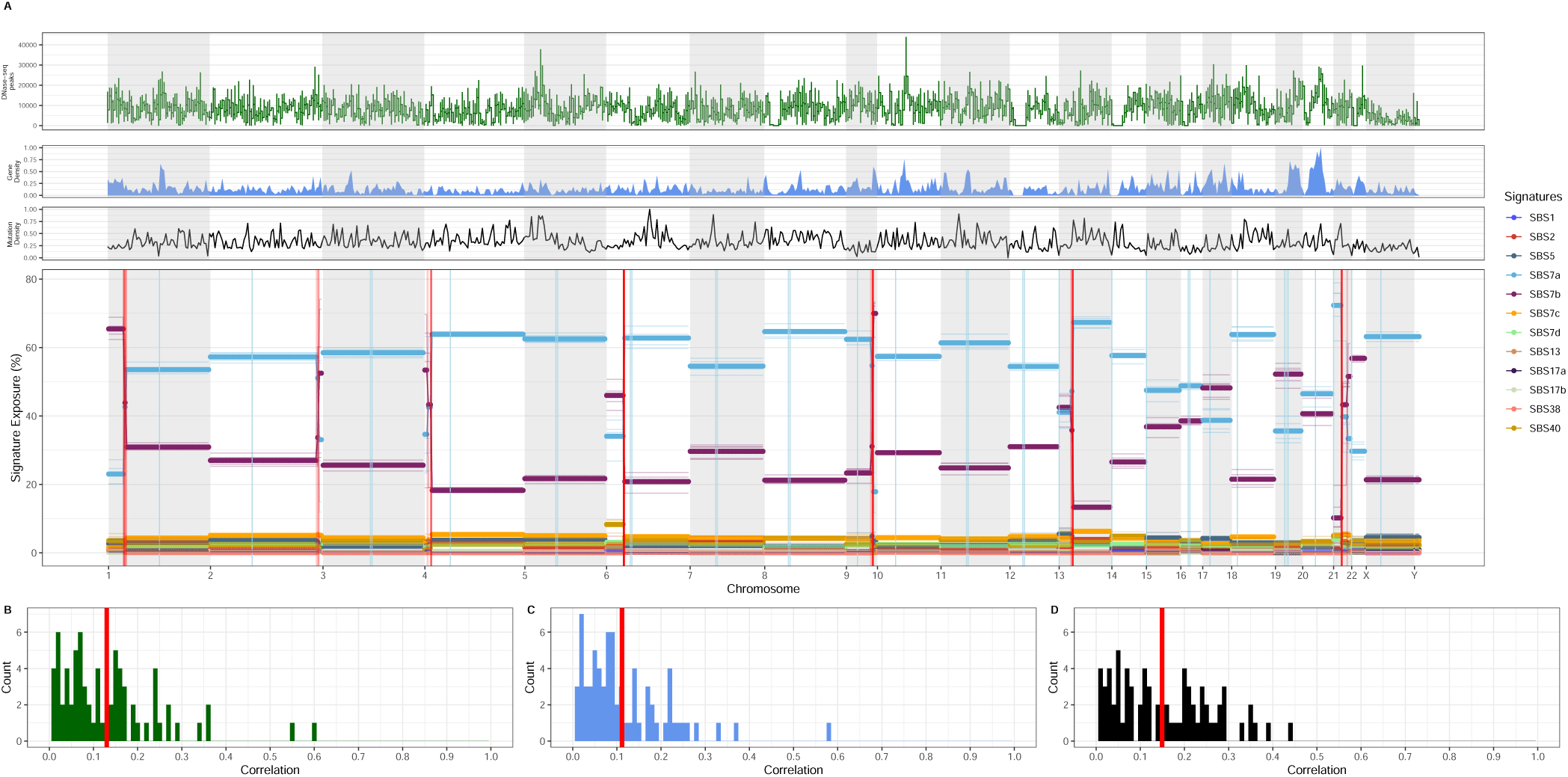
The association of signature activity profiles and chromatin accessibility, gene density, and mutation density profiles. A: Signature activity profile (chromosome-wise) for a representative melanoma sample with 168,604 mutations and 13 changepoints. Colored lines denote signature activities across bins of 200 mutations. Alternating gray and white bars denote chromosomal boundaries, and vertical blue lines show centromere positions. Red vertical lines show changepoint locations, and the opacity of these lines denotes confidence in that changepoint’s location. Above, black line plot depicts mutation density at each bin across the genome. Mutation density is normalized such that the bin with the highest density throughout the genome is scaled to one and the bin with the minimum density is scaled to zero. Above, blue area plot represents the average gene density at each bin, as determined from gene counts in hg19. Gene density is normalized in the same manner as mutation density. Above, green line plot depicts chromatin accessibility throughout a primary melanocyte genome as determined via DNAse-I accessibility index (Polak et al., 2015). Horizontal lines at each bin show the average chromatin accessibility across each bin, and vertical lines depict the range of chromatin accessibility values within each bin. B, C, and D show the distribution of correlation values between changepoint locations and DNAse-I accessibility index, gene density, and mutation density, respectively, across the 74 melanoma samples which contain changepoints. Mean correlation is highlighted in red on each plot.

To further investigate if changes in mutation density affect changepoint placement, we also analyzed the distances between changepoints and copy number aberrations (CNAs), and the distances between changepoints and kataegis events. We conducted a randomization test to determine, for each sample, if a significant proportion of the changepoints in the sample overlap with a CNA. We found that overall, 22/426 samples with changepoints have a higher proportion of changepoints overlapping with a CNA than would be expected at random (Table S2). Cervical cancers have the highest degree of changepoint/CNA co-occurrence, with three of 20 samples displaying significant overlap. No Melanoma, Lung-SCC, Colorect-AdenoCA, Lung-AdenoCA, Head-SCC, Stomach-AdenoCA, Prost-AdenoCA, Lymph-CLL, or Thy-AdenoCA samples showed significant changepoint/CNA overlap. These results emphasize that the changepoints detected by GenomeTrackSig reflect genuine shifts in mutational composition, not just mutation density. Further, we find that structural variation in the genome cannot explain the observed distribution of mutational signature activity changes.

Across the 2044/3059 changepoints within samples containing a kataegis event (Dentro et al., 2021, Nik-Zainal et al., 2012), these events were often distantly separated. The mean and median distances of changepoints to the nearest kataegis event were 214 Mb and 32 Mb, respectively, although some are relatively close: 357/2044 changepoints were located within 1 Mb from a kataegis event. These results are consistent with kataegis being one cause of a changepoint but not their primary cause. We observed a similar pattern when analyzing recurrent changepoints. The mean and median distances to a kataegis event among recurrent changepoints were 201 Mb and 18 Mb respectively, and 173/783 changepoints were located >1 Mb away from their nearest kataegis event. One reason for the partial association of kataegis and change-points may be that the vast majority of kataegis events involve APOBEC deaminases, which are represented by SBS2 and SBS13 (Alexandrov et al., 2020, Nik-Zainal et al., 2012). In this case, the hypermutation would result in a significant change in the local activity of these signatures, which would be detected by GenomeTrackSig. The localized hypermutation of the IGH locus in Lymph-BNHL may result in a SBS9-enriched kataegic event, which may explain both the high incidence of kataegis in Lymph-BNHL as well as the proximity of kataegis to the recurrent changepoints in that cancer (mean and median distances = 7 Mb and 6 Mb, respectively).

Lymph-BNHL exhibits at least one polymerase eta-driven hypermutation hotspot on every chromosome (Campbell et al., 2020), however changepoints were almost exclusively found on chromosome 14 (Figure 2). In contrast, Lymph-CLL contains eight recurrent changepoints across different chromosomes (Figure 2) but relatively few kataegis foci (Campbell et al., 2020). Although these two cancers have commonalities in their activities of SBS1, SBS9, SBS5, and SBS40 (Alexandrov et al., 2020) as well as being the only two cancer types known to display polymerase eta-driven kataegis (Campbell et al., 2020), within them we observe nearly opposite relationships between GenomeTrackSig changepoints and kataegis foci. In summary, kataegis has some association with GenomeTrackSig changepoints but it does not explain the majority of changepoints and its presence does not always give rise to a changepoint.

## Related Work

Wojtowicz et al. (2019) introduced SigMa, a composite multinomial mixture model and Hidden Markov Model to assign each mutation to a mutational signature. Transition probabilities between signatures are considered for adjacent clustered mutations, which are predominantly generated by kataegis events. GenomeTrackSig expands the number of regions which can be queried for a change in mutational signature activity to include regions of non-clustered mutations (Wojtowicz et al. (2019)’s “sky”mutations that are more than 2 kb from the next nearest mutation). However, transition points found by SigMa are not easily comparable with GenomeTrackSig changepoints, as a transition between generating processes may occur between individual mutations without necessarily accompanying a change in the relative activities of these mutational processes in the surrounding region.

## Discussion

Here we have introduced a new method, GenomeTrackSig, to detect region-specific changes in mutational signature activity within cancer genomes. Using this method we have demonstrated frequent changes in mutational signature activities over large chromosomal domains in a variety of cancers. We have also found a surprising number of recurrent changepoints in signature activities shared across cancers of the same type and among cancers of different types.

The scale, >1 Mb, at which GenomeTrackSig can generally detect changes in mutational signature activity is larger than regional factors already known to influence mutation rates, which makes it interesting that we see so many changepoints. For example, regional mutation rate varies due to changes in replication timing, differential activity of DNA repair mechanisms, and chromatin accessibility (Schuster-Böckler and Lehner, 2012, Supek and Lehner, 2019, Vöhringer et al., 2021, Yaacov et al., 2021, Supek and Lehner, 2015, Zheng et al., 2014). In samples with many mutations, in which a changepoint can demarcate smaller regions (<= 1 Mb in size), we may be able to attribute changes in signature activities to some of these factors that cause regional differences in the somatic mutation rate, especially if these factors change substantially over tumor development (see below). However, our effort to quantify this phenomenon using readily-measured genomic features – gene density, mutation density, structural and copy number variants, or kataegis (Table S2, Figure 5) – established that the signature changes we are detecting reflect a genuine shift in underlying mutational distributions and cannot be fully explained by these local, smaller-scale factors alone (Figure 5).

Perhaps, instead our changepoints demarcate large-scale chromosomal organization, with each large domain having its own underlying mutational distribution driven by domain-specific DNA damage and repair dynamics. Given that signature changes at recurrent changepoints are often consistent within a tissue yet variable across tissues, it is possible that such chromosomal domains can occur at similar locations in multiple tissue types yet exert tissue-specific effects on the mutational landscape. One intriguing possibility is that these large-scale domains are delineated by the 3D organization of the cancer genome – representing, for example, larger topologically associating domains (TADs) Yu and Ren (2017).

The signature dynamics at the most common changepoint in melanoma samples support our hypothesis that change-points are driven by chromatin state variation over large scales. The changepoint region at 1:47,000,000-51,000,000 is found in 63 samples from two tumor types, 55 of those melanomas. As shown in Figure 3, all 55 melanomas display strikingly consistent decreases in SBS7b activity and increases in SBS7a activity at this changepoint. An analysis of the genomic properties influencing mutational signature activities by Vöhringer et al. (2021), using a *de novo* signature extraction method, identified two signatures exclusively occurring in skin cancers which highly resemble SBS7a and SBS7b in terms of their mutational distributions and associations with UV exposure. Their study found that the SBS7a-like signature has high activity in quiescent chromatin, whereas SBS7b-like is enriched in active chromatin and has a strong transcriptional strand bias. They suggested this differential signature activity across chromatin states reflects different operative DNA repair mechanisms. In this scenario, SBS7a-like activity reflects UV damage cleared by global genome nucleotide excision repair (GG-NER), which operates in quiescent and active regions, and SBS7b-like activity reflects damage cleared by a combination of GG-NER and transcription-coupled nucleotide excision repair (TC-NER), the latter of which operates in open chromatin and is activated by template strand lesions on actively transcribed genes (Vöhringer et al., 2021). Therefore, the activity shifts between SBS7a and SBS7b that we observe at recurrent changepoint regions in melanomas may reflect large-scale changes in chromatin state, coupled with changes in active DNA repair processes.

Based on signature activity profiles from Lymph-CLL samples, we also hypothesize that large-scale changes in chromatin state sometimes occur in a timing-dependent manner. We observe changepoints in which early-development signatures decrease in activity and late-development signatures increase in activity, and vice versa. Cancer development is marked by a redistribution of the mutational landscape, as exposures to mutagens and DNA repair failures strip active genes in euchromatic regions of the preferential repair that they normally receive (Supek and Lehner, 2019). Thus, we may see higher activity of late-development signatures in regions that normally have a low mutation rate, but which become vulnerable to mutation during subclonal expansion when different mutational processes are active than were early on. Conversely, late-replicating regions of closed chromatin might display higher activity of early-development signatures since these tend to have higher mutation rates under normal conditions.

Via mutational signature analysis, we have identified that mutational signature activities change over chromosomal domains. These changes can be highly consistent on multiple levels. We hypothesize that, among other factors, our method is detecting the impacts of large-scale regional changes in chromatin state. Recurrent changepoints in melanoma samples (Figure 2) provide compelling evidence that regions on either side of a changepoint differ in terms of their chromatin accessibility and DNA repair dynamics (Vöhringer et al., 2021). Furthermore, changepoints that appear in multiple tissue types raise the possibility that such wide-scale epigenomic changes are common events in the course of cancer development. These results call for further exploration into how factors like chromatin state and DNA repair mechanisms vary over wider domains, and which elements of these changes are characteristic of many cancers and which are tissue-specific. Interestingly, changepoints sometimes reflect a shift in activity between signatures characteristic of early tumor development and signatures characteristic of subclonal expansion. Thus, we hypothesize that large-scale changes in chromatin accessibility may also occur in a timing-dependent manner, and can demarcate regions that are more likely to be impacted by mutations early or late in a cancer’s evolutionary trajectory. This finding may be clinically relevant given that intra-tumor heterogeneity is a mechanism of therapeutic resistance and therefore presents significant challenges for treatment (Maley et al., 2006, McGranahan and Swanton, 2017, Mroz et al., 2013). Therefore, genome-wide mutational signature analysis can help us further characterize and localize the genomic and epigenomic changes that occur during tumor development.

## ACKNOWLEDGEMENTS

This research was supported by an NIH/NCI Cancer Center Support Grant P30 CA008748 and an NCI R25 CA233208 grant. Q.M. is a CCAI CIFAR chair. C.T. was supported by the Tri-I Computational Biology Summer Program.

## Supplementary Note 1: Supplementary

**Table S1.**
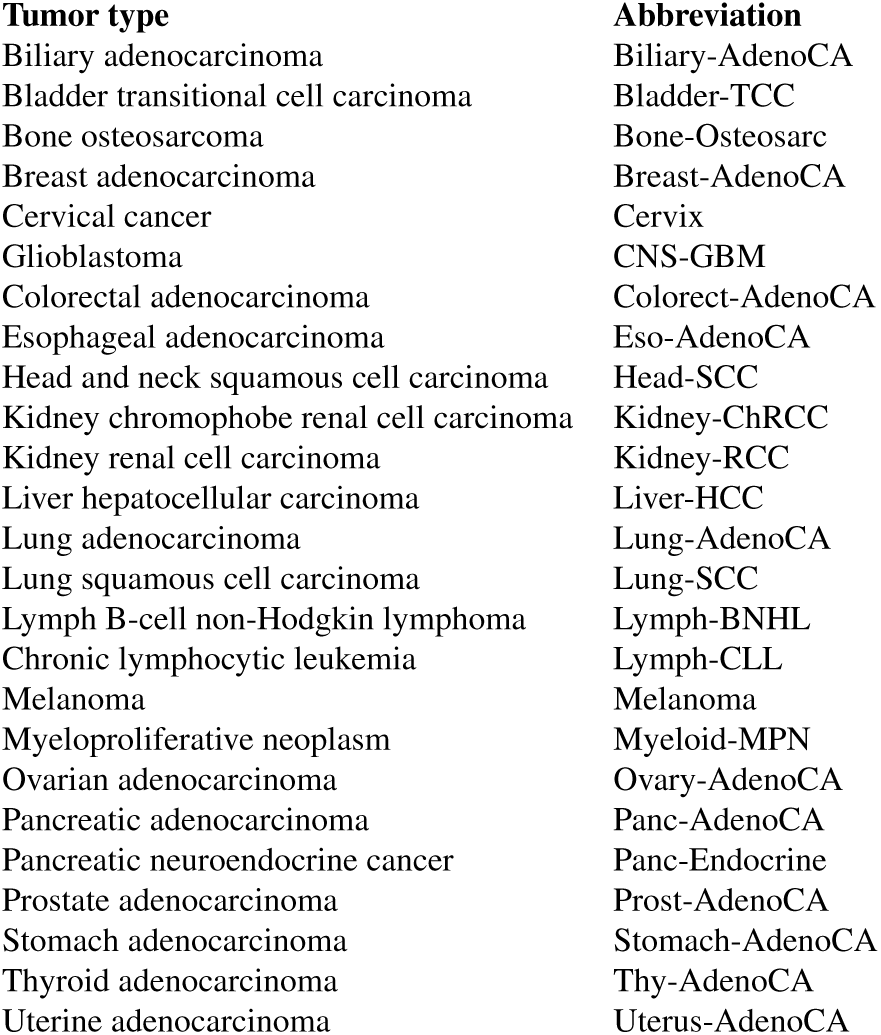
Abbreviations of tumor types analyzed in the study.

**Table S2.**
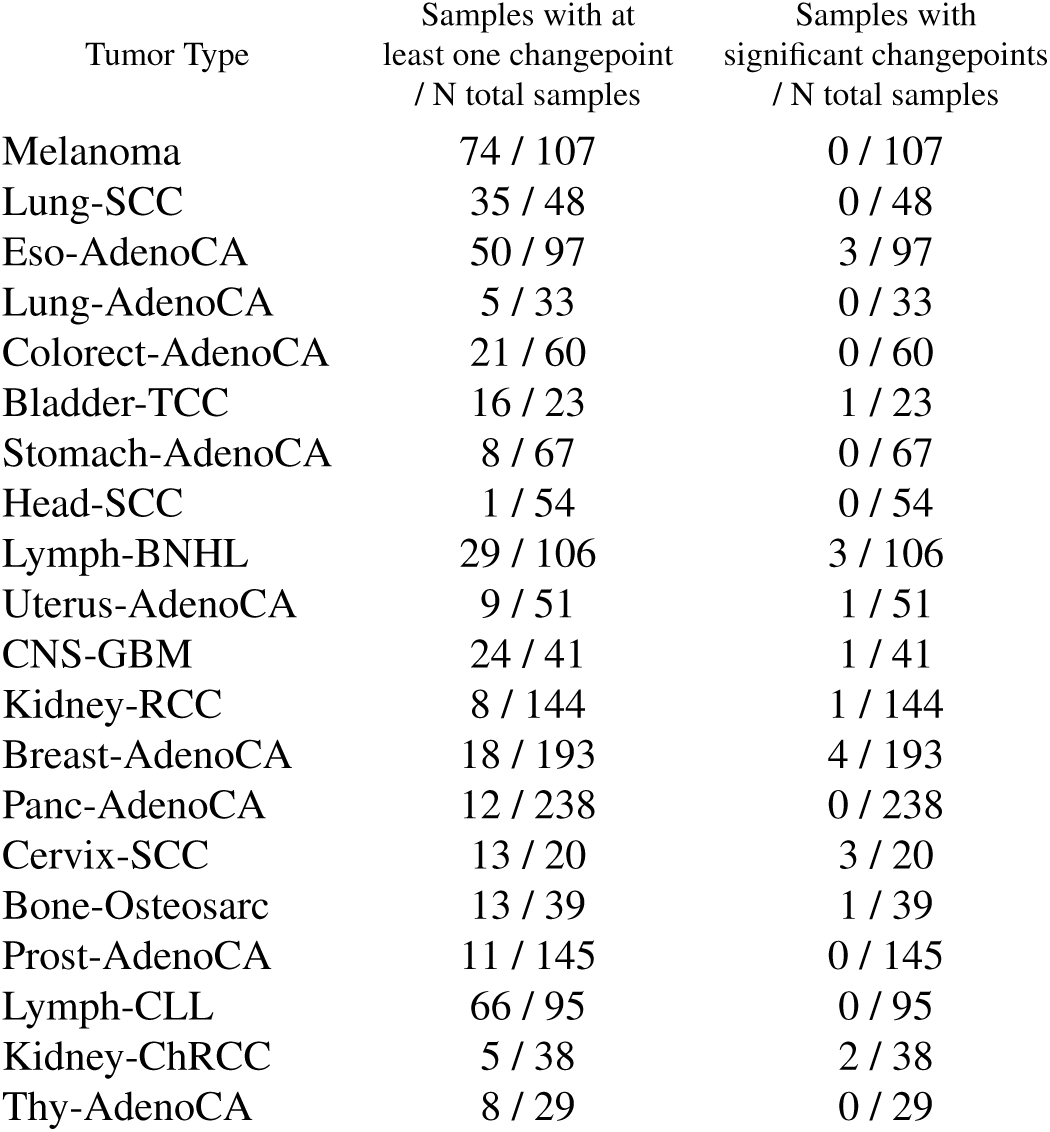
Co-occurrence of changes in mutational signature activities and copy number aberrations across 20 tumor types. For each tumor type, we conducted a randomization test to determine how many samples have a significant proportion of their changepoints overlapping with a copy number aberration (CNA). Randomization tests were conducted by generating 10,000 random samples with the same number of changepoints and same changepoint region spans as the original samples with changepoint locations randomized. Each random set of change-points was compared to the original sample copy number profile to calculate the proportion of changepoints overlapping with a CNA. This set of proportions formed the null distribution against which the sample proportion was compared to determine a p-value (*α*=0.05). P-values were adjusted for multiple comparisons with Bonferroni corrections.

| Tumor Type | Samples with at least one changepoint / N total samples | Samples with significant changepoints / N total samples |
| --- | --- | --- |
| Melanoma | 74 / 107 | 0 / 107 |
| Lung-SCC | 35 / 48 | 0 / 48 |
| Eso-AdenoCA | 50 / 97 | 3 / 97 |
| Lung-AdenoCA | 5 / 33 | 0 / 33 |
| Colorect-AdenoCA | 21 / 60 | 0 / 60 |
| Bladder-TCC | 16 / 23 | 1 / 23 |
| Stomach-AdenoCA | 8 / 67 | 0 / 67 |
| Head-SCC | 1 / 54 | 0 / 54 |
| Lymph-BNHL | 29 / 106 | 3 / 106 |
| Uterus-AdenoCA | 9 / 51 | 1 / 51 |
| CNS-GBM | 24 / 41 | 1 / 41 |
| Kidney-RCC | 8 / 144 | 1 / 144 |
| Breast-AdenoCA | 18 / 193 | 4 / 193 |
| Panc-AdenoCA | 12 / 238 | 0 / 238 |
| Cervix-SCC | 13 / 20 | 3 / 20 |
| Bone-Osteosarc | 13 / 39 | 1 / 39 |
| Prost-AdenoCA | 11 / 145 | 0 / 145 |
| Lymph-CLL | 66 / 95 | 0 / 95 |
| Kidney-ChRCC | 5 / 38 | 2 / 38 |
| Thy-AdenoCA | 8 / 29 | 0 / 29 |

**Fig. S1.**
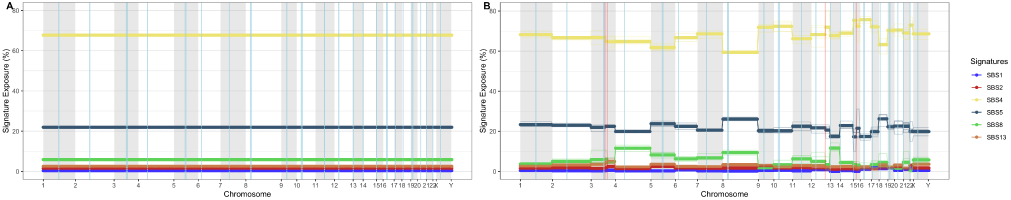
Comparison of genome-wise (A) and chromosome-wise (B) activity profiles constructed by GenomeTrackSig. Input data is a Lung-SCC genome with 78,839 mutations. A bin size of 200 mutations was used and 5 bootstraps were performed for each experiment. Each point is a signature activity estimate at one bin of mutations. Alternating gray and white bars distinguish chromosomes and blue vertical lines show centromere positions. Red vertical lines denote changepoints, and the opacity of changepoints represents their bootstrap support.

**Fig. S2.**
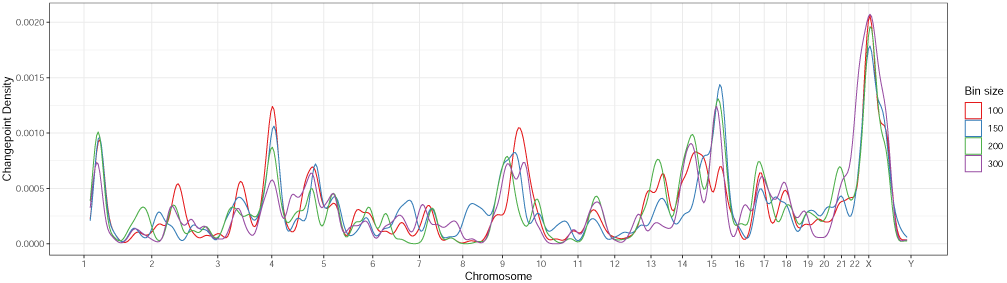
Lung-SCC changepoint density across the genome at four different bin sizes. GenomeTrackSig was run genome-wise on 32 Lung-SCC samples with a bin size of either 100, 150, 200, or 300. Density plot across the genome of pooled changepoint positions in all samples is shown for each bin size analyzed.

**Fig. S3.**
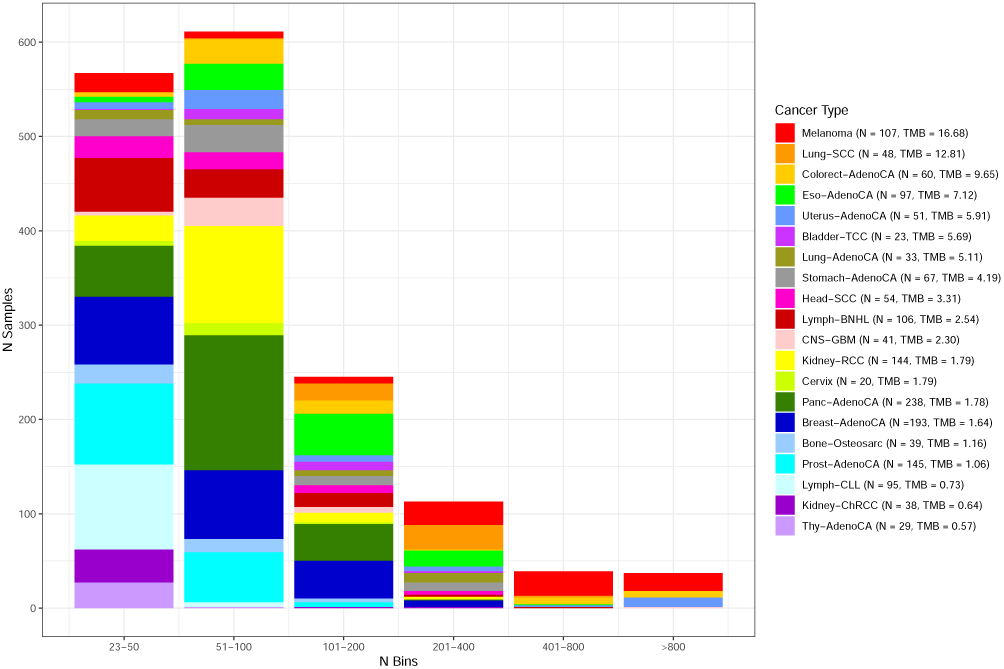
Number of bins per sample across cancer types. Stacked bar chart depicting the distribution of bin numbers across samples, colored by cancer type. Minimum number of bins is 23 (Melanoma, Lung-AdenoCA, Lymph-BNHL, Kidney-RCC, Prost-AdenoCA, Lymph-CLL, Kidney-ChRCC, Thy-AdenoCA) and maximum number of bins is 2,895 (Colorect-AdenoCA). Sample size and geometric mean TMB are shown for each cancer type.

**Fig. S4.**
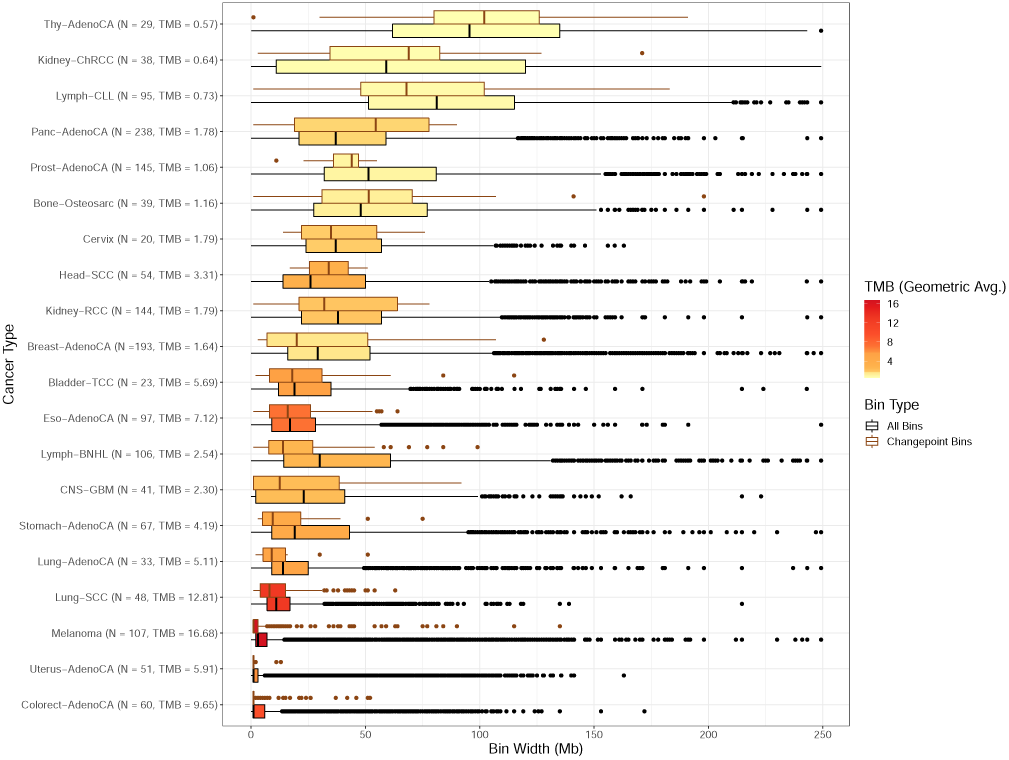
Distribution of bin widths by cancer type. Boxplots showing the range of bin widths, in megabases, for all samples analyzed in the study. Boxplots are outlined according to the type of bin plotted, either all bins or bins containing change-points. Sample size and geometric mean TMB are shown for each cancer type. Boxplots are colored according to geometric mean TMB.

**Fig. S5.**
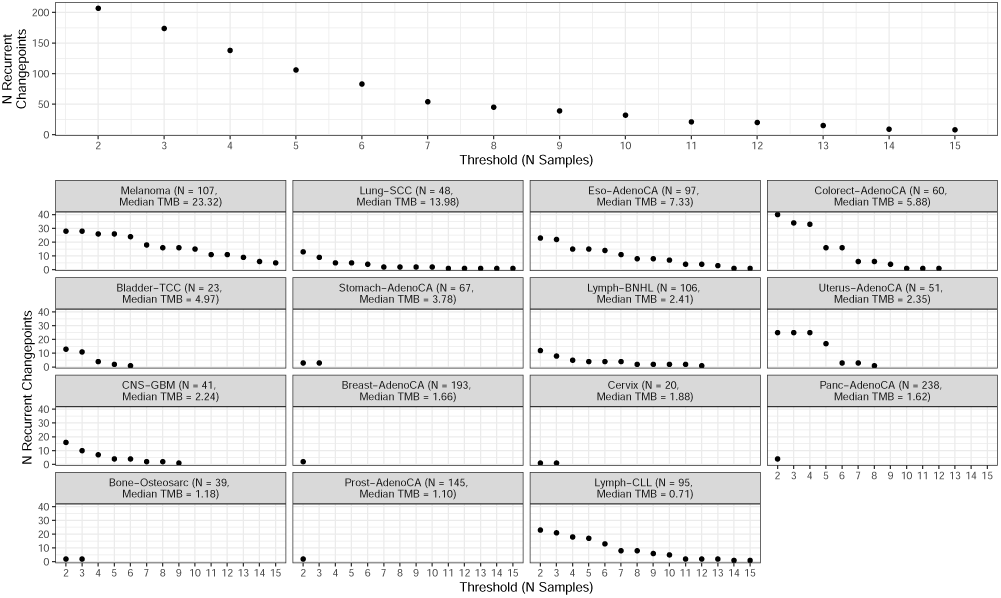
Number of recurrent changepoints identified at different sample thresholds. Top: Number of recurrent changepoints identified across all cancer types depending on which number of samples is used as the threshold to determine which changepoints are considered ‘recurrent.’ Bottom: Number of recurrent changepoints identified in each cancer type compared to the sample threshold. Sample size and median tumor mutational burden are shown for each cancer type.

