## Supplementary Note 1 for "Regional Mutational Signature Activities in Cancer Genomes"

**Table S1:** Abbreviations of tumor types analyzed in the study.

| Tumor type | Abbreviation |
|---|---|
| Biliary adenocarcinoma | Biliary-AdenoCA |
| Bladder transitional cell carcinoma | Bladder-TCC |
| Bone osteosarcoma | Bone-Osteosarc |
| Breast adenocarcinoma | Breast-AdenoCA |
| Cervical cancer | Cervix |
| Glioblastoma | CNS-GBM |
| Colorectal adenocarcinoma | Colorect-AdenoCA |
| Esophageal adenocarcinoma | Eso-AdenoCA |
| Head and neck squamous cell carcinoma | Head-SCC |
| Kidney chromophobe renal cell carcinoma | Kidney-ChRCC |
| Kidney renal cell carcinoma | Kidney-RCC |
| Liver hepatocellular carcinoma | Liver-HCC |
| Lung adenocarcinoma | Lung-AdenoCA |
| Lung squamous cell carcinoma | Lung-SCC |
| Lymph B-cell non-Hodgkin lymphoma | Lymph-BNHL |
| Chronic lymphocytic leukemia | Lymph-CLL |
| Melanoma | Melanoma |
| Myeloproliferative neoplasm | Myeloid-MPN |
| Ovarian adenocarcinoma | Ovary-AdenoCA |
| Pancreatic adenocarcinoma | Panc-AdenoCA |
| Pancreatic neuroendocrine cancer | Panc-Endocrine |
| Prostate adenocarcinoma | Prost-AdenoCA |
| Stomach adenocarcinoma | Stomach-AdenoCA |
| Thyroid adenocarcinoma | Thy-AdenoCA |

| | |
|---|---|
| Uterine adenocarcinoma | Uterus-AdenoCA |

**Table S2: Co-occurrence of changes in mutational signature activities and copy number aberrations across 20 tumor types.** For each tumor type, we conducted a randomization test to determine how many samples have a significant proportion of their changepoints overlapping with a copy number aberration (CNA). Randomization tests were conducted by generating 10,000 random samples with the same number of changepoints and same changepoint region spans as the original samples with changepoint locations randomized. Each random set of changepoints was compared to the original sample copy number profile to calculate the proportion of changepoints overlapping with a CNA. This set of proportions formed the null distribution against which the sample proportion was compared to determine a p-value ($\alpha = 0.05$). P-values were adjusted for multiple comparisons with Bonferroni corrections.

| Tumor Type | Samples with at least one changepoint / N total samples | Samples with significant changepoint / n samples with at least one changepoint |
|---|---|---|
| Melanoma | 74 / 107 | 0 / 107 |
| Lung-SCC | 35 / 48 | 0 / 48 |
| Eso-AdenoCA | 50 / 97 | 3 / 97 |
| Lung-AdenoCA | 5 / 33 | 0 / 33 |
| Colorect-AdenoCA | 21 / 60 | 0 / 60 |
| Bladder-TCC | 16 / 23 | 1 / 23 |
| Stomach-AdenoCA | 8 / 67 | 0 / 67 |
| Head-SCC | 1 / 54 | 0 / 54 |
| Lymph-BNHL | 29 / 106 | 3 / 106 |
| Uterus-AdenoCA | 9 / 51 | 1 / 51 |
| CNS-GBM | 24 / 41 | 1 / 41 |
| Kidney-RCC | 8 / 144 | 1 / 144 |
| Breast-AdenoCA | 18 / 193 | 4 / 193 |
| Panc-AdenoCA | 12 / 238 | 2 / 238 |
| Bone-Osteosarc | 13 / 39 | 1 / 39 |
| Prost-AdenoCA | 11 / 145 | 0 / 145 |

| | | |
|---|---|---|
| Lymph-CLL | 66 / 95 | 0 / 95 |
| Kidney-ChRCC | 5 / 38 | 2 / 38 |
| Cervix-SCC | 13 / 20 | 3 / 20 |
| Thy-AdenoCA | 8 / 29 | 0 / 29 |

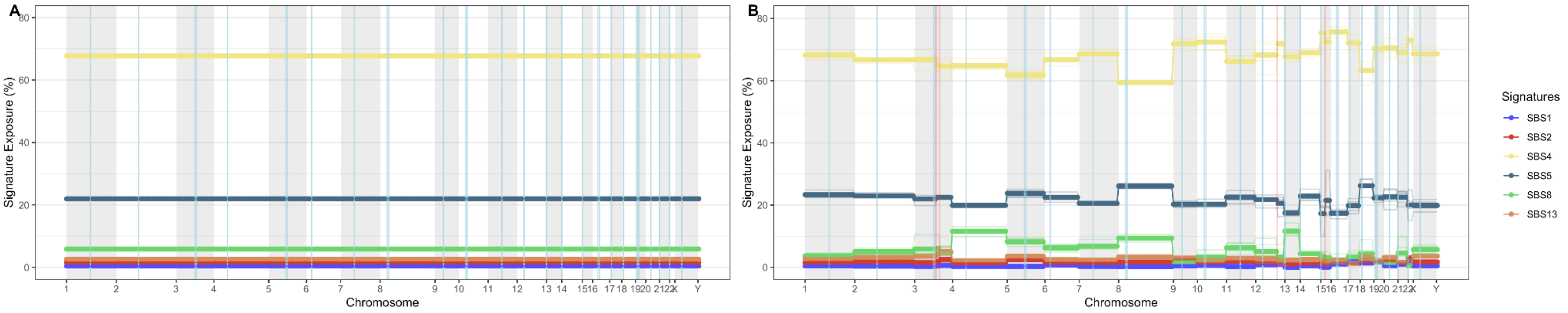


**Figure S1: Comparison of genome-wise (A) and chromosome-wise (B) activity profiles constructed by TrackSig.** Input data is a Lung-SCC genome with 78,839 mutations. A bin size of 200 mutations was used and 5 bootstraps were performed for each experiment. Each point is a signature activity estimate at one bin of mutations. Alternating gray and white bars distinguish chromosomes and blue vertical lines show centromere positions. Red vertical lines denote changepoints, and the opacity of changepoints represents their bootstrap support.

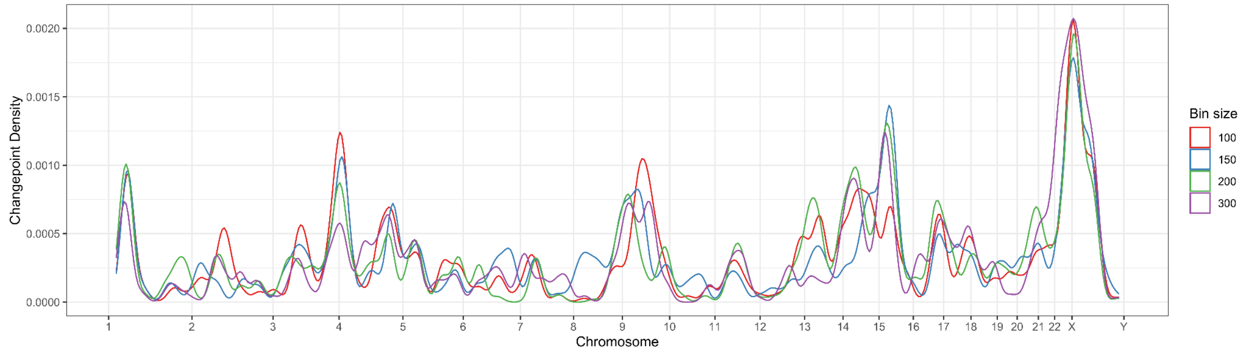


**Figure S2: Lung-SCC changepoint density across the genome at four different bin sizes.** GenomeTrackSig was run genome-wise on 32 Lung-SCC samples with a bin size of either 100, 150, 200, or 300. Density plot across the genome of pooled changepoint positions in all samples is shown for each bin size analyzed.

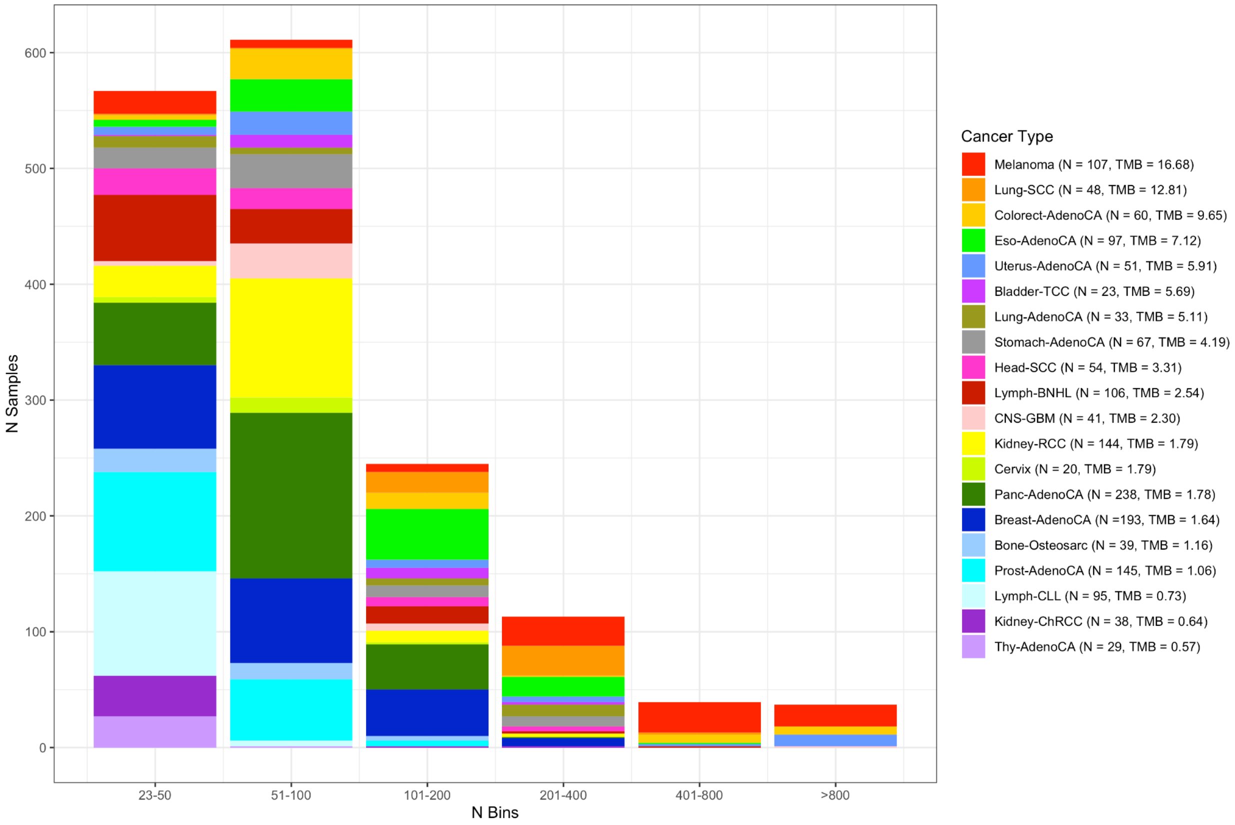


**Figure S3: Number of bins per sample across cancer types.** Stacked bar chart depicting the distribution of bin numbers across samples, colored by cancer type. Minimum number of bins is 23 (Melanoma, Lung-AdenoCA, Lymph-BNHL, Kidney-RCC, Prost-AdenoCA, Lymph-CLL, Kidney-ChRCC, Thy-AdenoCA) and maximum number of bins is 2895 (Colorect-AdenoCA). Sample size and geometric mean TMB are shown for each cancer type.

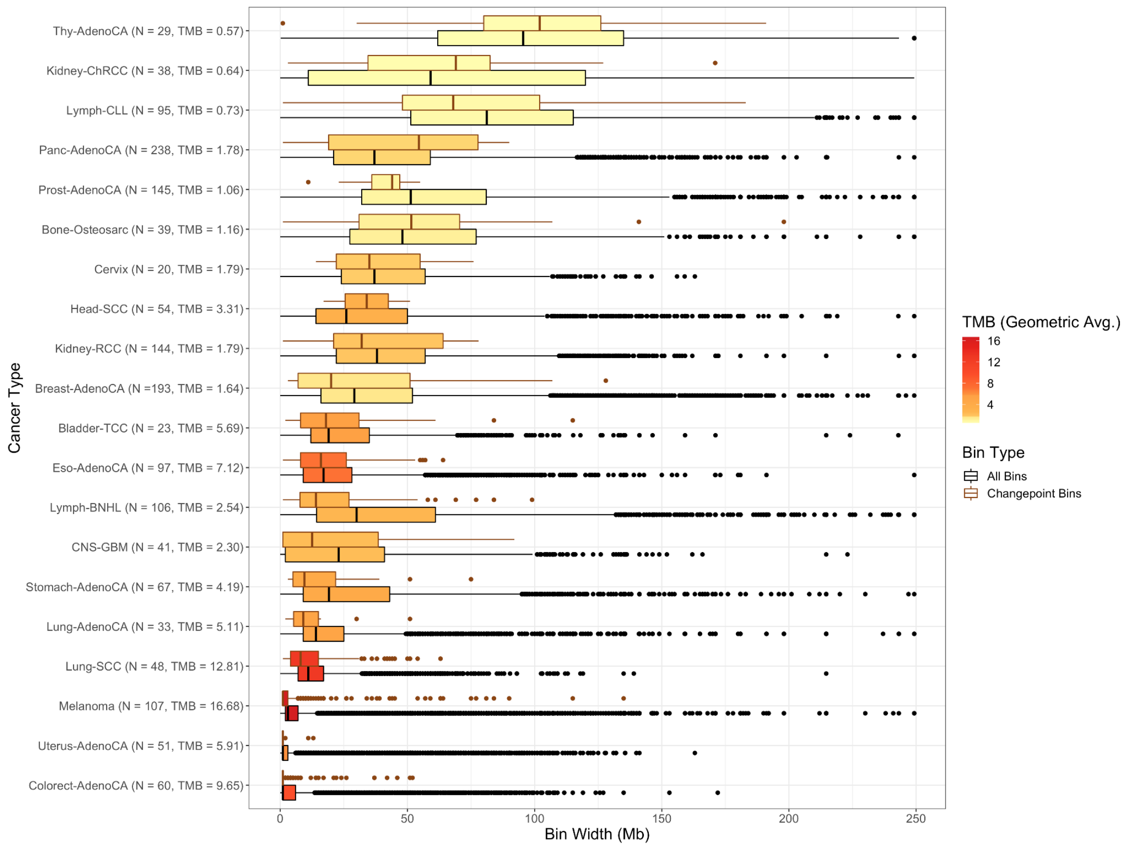


**Figure S4: Distribution of bin widths by cancer type.** Boxplots showing the range of bin widths, in megabases, for all samples analyzed in the study. Boxplots are outlined according to the type of bin plotted, either all bins or bins containing changepoints. Sample size and geometric mean TMB are shown for each cancer type. Boxplots are colored according to geometric mean TMB.

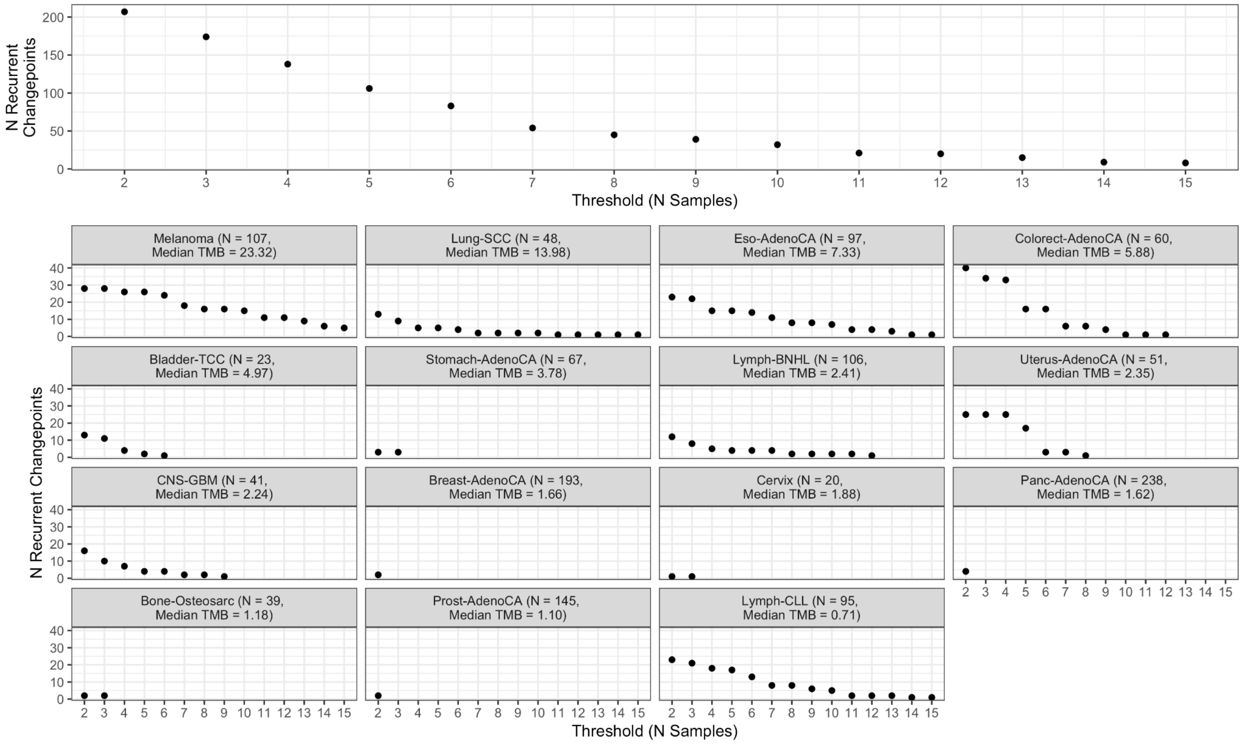


**Figure S5: Number of recurrent changepoints identified at different sample thresholds.** Top: Number of recurrent changepoints identified across all cancer types depending on which number of samples is used as the threshold to determine which changepoints are considered 'recurrent.' Bottom: Number of recurrent changepoints identified in each cancer type compared to the sample threshold. Sample size and median tumor mutational burden is shown for each cancer type.
